# Overexpression of Ribosomal Proteins Leads to Zn Resistance in *Escherichia coli*

**DOI:** 10.1101/2024.10.02.616313

**Authors:** Tomoki Kosaki, Riko Shirakawa, Kazuya Ishikawa, Kazuyuki Furuta, Chikara Kaito

## Abstract

Knockout of ribosomal protein bL36 (RpmJ) leads to Zn resistance in *Escherichia coli*, and the expression of ribosomal protein genes other than RpmJ increase in the *rpmJ*-knockout strain. In this study, we examined whether the overexpression of ribosomal proteins causes Zn resistance using an *E. coli* overexpression gene library (ASKA clone library). The overexpression of 48 of the 54 ribosomal proteins led to Zn resistance. However, the overexpression of proteins other than ribosomal proteins did not lead to Zn resistance, suggesting that Zn resistance is a phenomenon specific to the overexpression of ribosomal proteins. In addition, the overexpression of ribosomal proteins did not lead to resistance to metal ions other than Zn (Cu^2+^, Ni^2+^, Mn^2+^, and Ag^+^), suggesting a Zn-specific resistance mechanism. Deletion of ZntA, a Zn efflux pump, resulted in the loss of Zn resistance in a ribosomal protein-overexpressing strain. Deletion of Lon protease, which is responsible for degrading misfolded proteins, in a ribosomal protein-overexpressing strain resulted in the accumulation of overexpressed ribosomal proteins and loss of Zn resistance. These results suggest that the overexpression of ribosomal proteins leads to Zn resistance in *E. coli via* ZntA and Lon protease.

**Importance:** The ribosome is a complex comprising ribosomal RNAs and more than 50 types of ribosomal proteins. Ribosomal proteins play an important role in ribosomal function responsible for protein translation; however, their involvement in other cellular processes is not fully understood. Based on the finding that ribosomal protein expression increases in a Zn-resistant *E. coli* mutant, we analyzed 54 ribosomal proteins and found that the overexpression of 48 ribosomal proteins led to Zn resistance. This finding suggests a role for ribosomal proteins in resistance to zinc stress.

## Introduction

Zn, a trace element, functions as a cofactor for several enzymes and transcription factors in the cell, and plays a pivotal role in bacterial survival and growth (1). Excess Zn is highly toxic to bacteria and interferes with bacterial metabolism by disrupting the iron-sulfur clusters of several enzymes, such as succinate dehydrogenase (2) and peptidoglycan biosynthetic enzymes (3), and disturbing metal homeostasis (4, 5). Taking advantage of the toxic properties of Zn, human immune macrophages expose bacteria to excess Zn to kill them (6). Bacteria utilize the Zn efflux pumps ZntA and ZitB to regulate intracellular Zn levels (7, 8, 9).

Ribosomal proteins play a central role in protein synthesis (10) and are involved in cellular stress responses (11, 12). For example, in *Escherichia coli*, under amino acid starvation conditions, free ribosomal proteins are preferentially degraded by Lon protease to supply amino acids (13). In *Bacillus subtilis*, ribosomal protein L11 is required for activating Sigma B, a transcription factor involved in diverse stress responses (14). In eukaryotes, when RNA synthesis is inhibited, several ribosomal proteins inhibit the activity of the ubiquitin ligase mouse double minute 2 homolog, thereby stabilizing its target, the tumor suppressor p53 (15).

We have previously noticed that knocking out several ribosomal proteins triggers Zn resistance in *E. coli* (16). Furthermore, the *rpmJ*-knockout mutant shows altered translation fidelity and increased expression of ribosomal proteins other than ribosomal protein bL36 (RpmJ) (16). Zn resistance in the *rpmJ*-knockout mutant is lost by knocking out ZntA (16). These findings suggest that the loss of RpmJ affects translation and leads to Zn resistance *via* ZntA; however, the molecular mechanism of Zn resistance remains unclear.

In this study, based on the finding of increased expression of ribosomal proteins in the *rpmJ*-knockout mutant, we tested whether overexpression of all 54 ribosomal proteins (17) causes ZntA-mediated Zn resistance in *E. coli*. Additionally, we explored the role of Lon protease in the Zn resistance caused by the ribosomal proteins.

## Results

### Overexpression of ribosomal proteins leads to Zn resistance in *E. coli*

First, we transformed BW25113 (WT) with the ASKA clone library to generate strains overexpressing 54 different ribosomal proteins and examined the sensitivity of the strain to Zn. In the absence of Zn, most strains overexpressing ribosomal proteins showed slightly lower CFU than did the empty vector-transformed strain, whereas in the presence of Zn, 48 of the 54 ribosomal protein-overexpressing strains showed higher CFU than did the empty vector-transformed strain (**Fig. 1**). These results suggest that overexpression of most ribosomal proteins leads to Zn resistance in *E. coli*.

**FIG 1.**
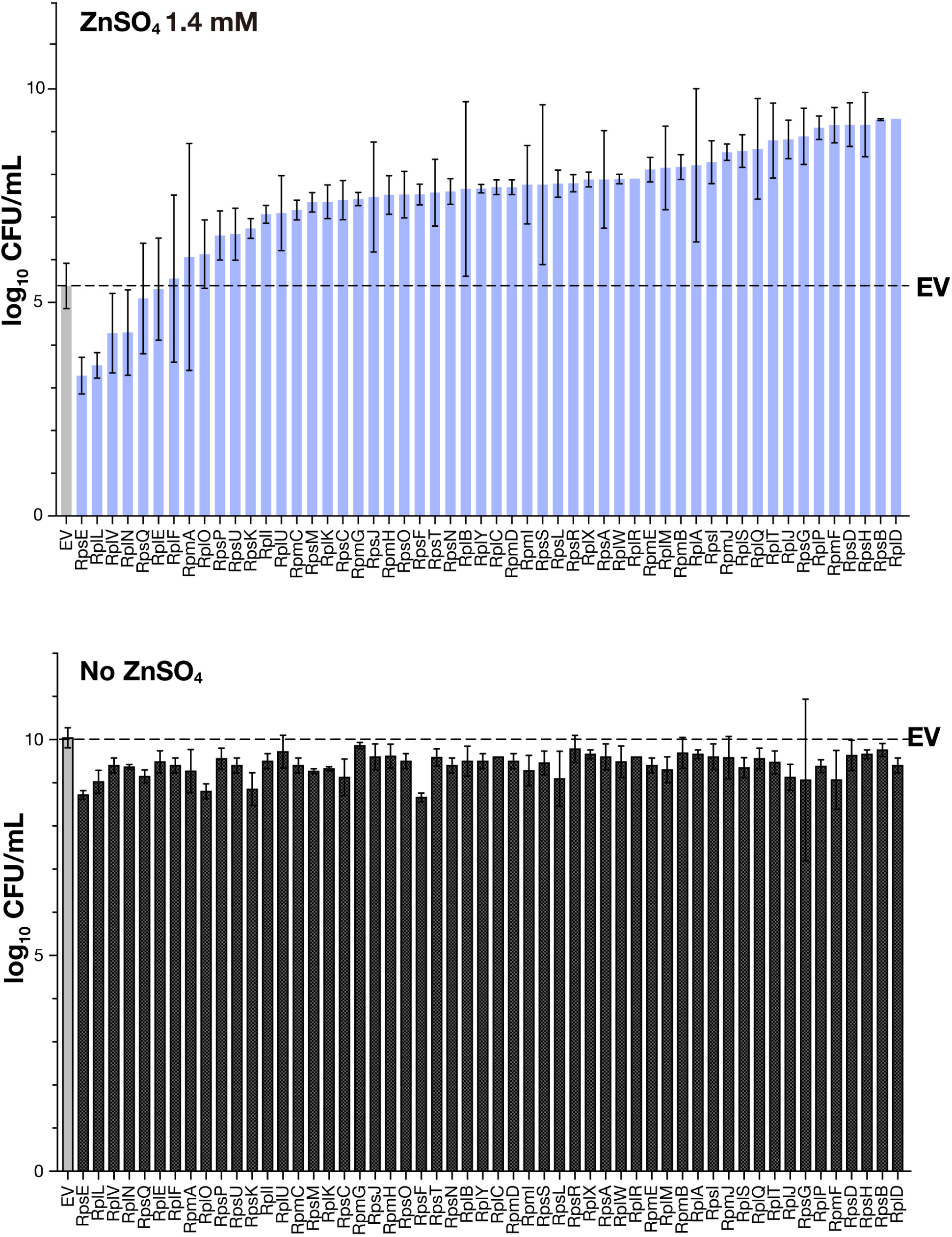
Overexpression of ribosomal proteins causes Zn resistance. Ten-fold serial dilutions of overnight cultures of an empty vector-transformed *E. coli* strain (WT/pCA24N) and 54 ribosomal protein-overexpressing *E. coli* strains were spotted on LB agar plates containing 0.1 mM IPTG with or without 1.4 mM Zn. After overnight incubation, colonies were counted. Data represent mean ± SD of three independent experiments.

### Overexpression of ribosomal proteins does not lead to resistance to Ag^+^, Ni^2+^, Mn^2+^, and Cu^2+^

Since the overexpression of ribosomal proteins led to zinc resistance, we examined whether the overexpression of ribosomal proteins could lead to resistance to metal ions other than zinc ions; Ag^+^, Ni^2+^, Mn^2+^, and Cu^2+^. Most ribosomal protein-overexpressing strains did not show resistance to Ag^+^, Ni^2+^, Mn^2+^, and Cu^2+^ (**Fig. S1, S2**). In addition, the correlation between the degree of resistance to these metal ions and degree of Zn resistance was low (r = 0.1617–0.3004) (**Fig. 2**). These results suggest that the overexpression of ribosomal proteins leads to Zn-specific resistance.

**FIG 2.**
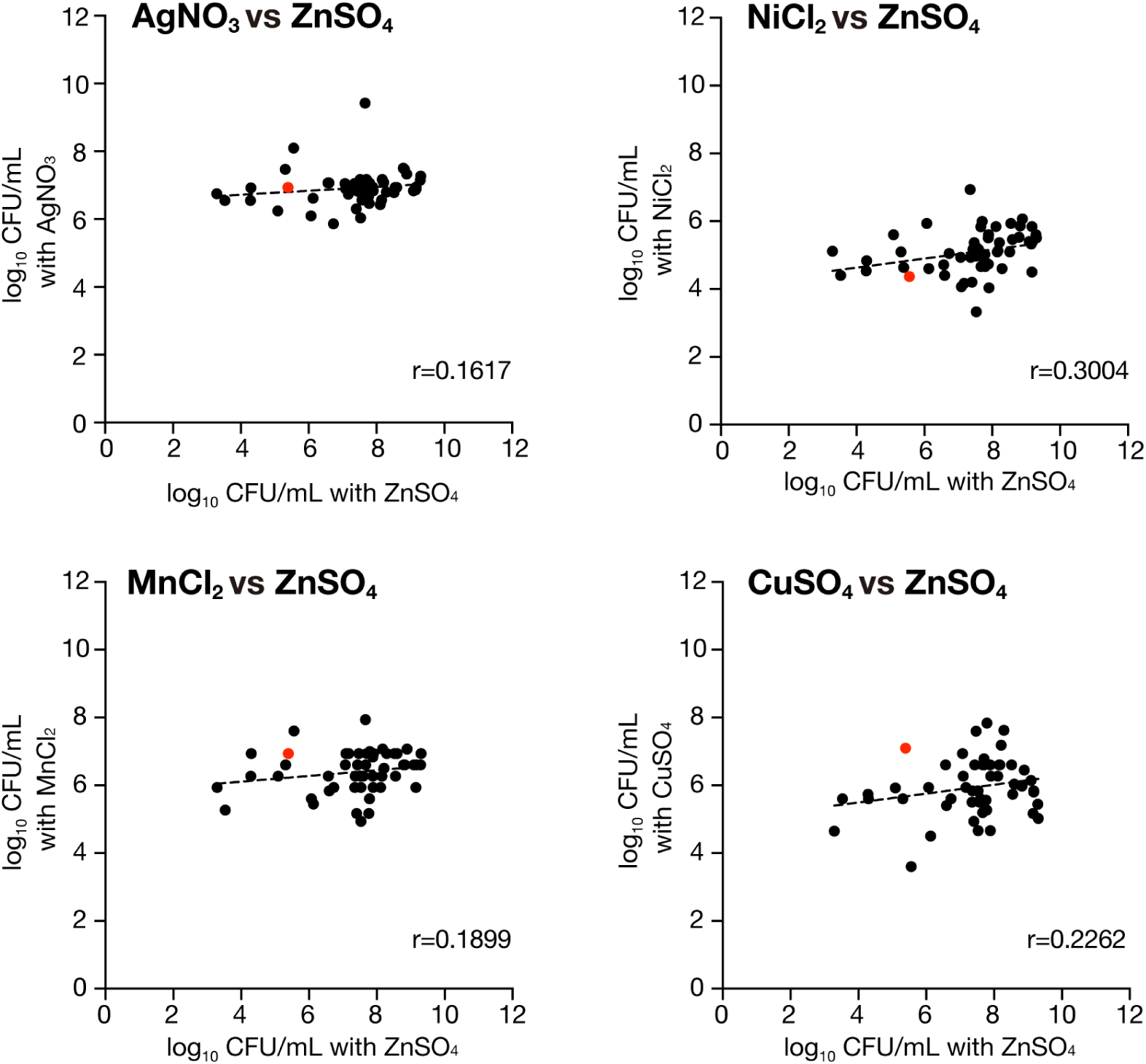
Correlation analyses between Zn resistance and other metal resistances of ribosomal protein-overexpressing strains. Ten-fold serial dilutions of overnight cultures of an empty vector-transformed *E. coli* strain (WT/pCA24N) and 54 ribosomal protein-overexpressing *E. coli* strains were spotted on LB agar plate containing 0.1 mM IPTG with Ag^+^, Ni^2+^, Mn^2+^, or Cu^2+^ ions, and colonies were counted after overnight incubation. The number of colonies are presented in Figures S1 and S2. Correlation between the number of colonies formed in presence of Zn and other metal ions were analyzed.

### The degree of Zn resistance of the strains overexpressing ribosomal proteins correlates with the degree of resistance to streptomycin

We examined whether the overexpression of ribosomal proteins leads to resistance to the ribosome-targeting antibiotics tetracycline, streptomycin, and erythromycin. Some ribosomal protein-overexpressing strains were found to be resistant to these antibiotics (**Fig. S3 and S4**). Furthermore, the correlation between the degree of resistance to erythromycin or tetracycline and degree of Zn resistance was low (r = 0.1468–0.2151), whereas a strong correlation was observed between the degree of resistance to streptomycin and degree of Zn resistance (r = 0.7172) (**Fig. 3)**. These results suggest a common mechanism between resistance to streptomycin and Zn in ribosomal protein-overexpressing strains.

**FIG 3.**
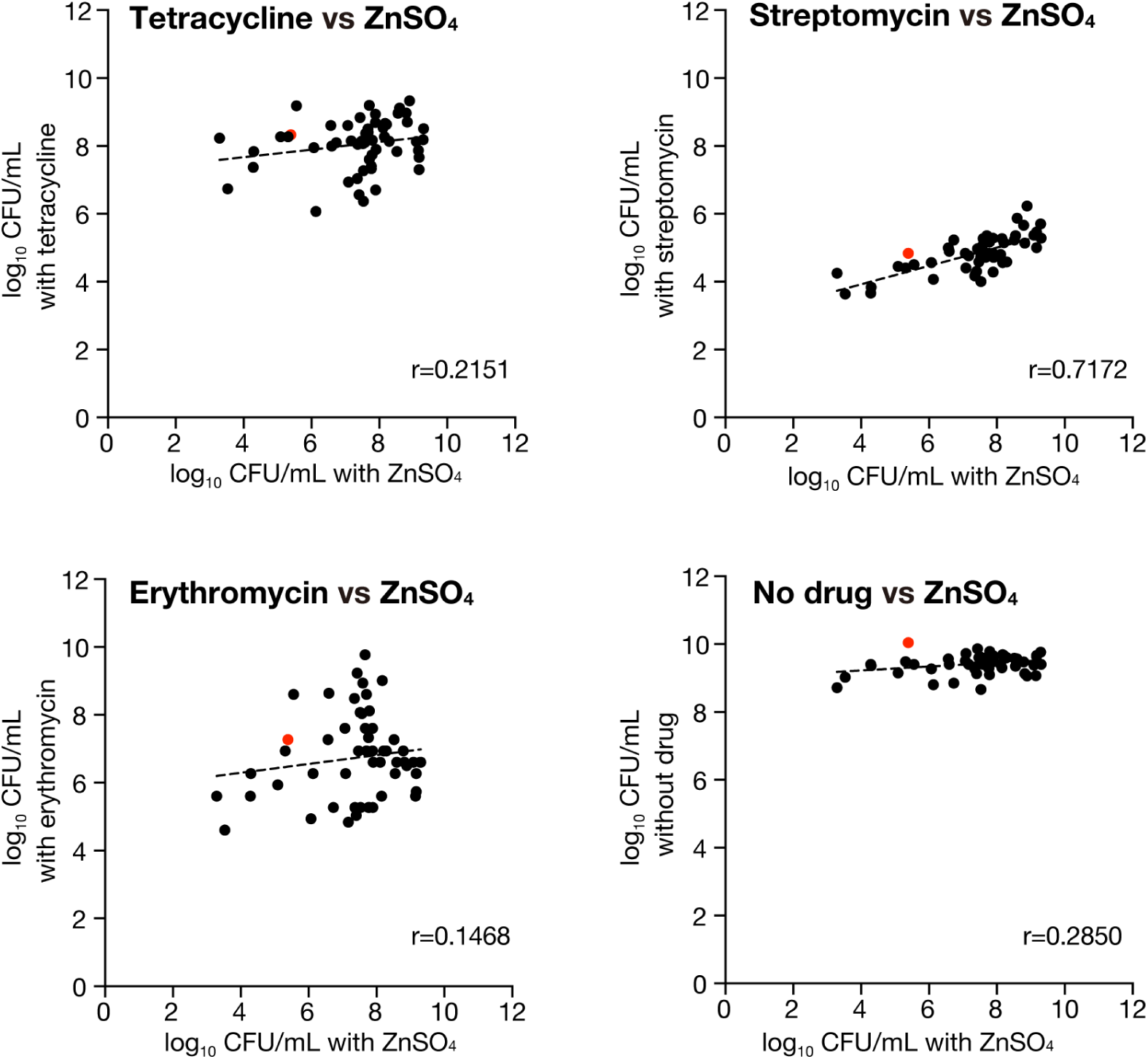
Correlation analyses between Zn resistance and antibiotic resistance of ribosomal protein-overexpressing strains. Ten-fold serial dilutions of overnight cultures of an empty vector-transformed *E. coli* strain (WT/pCA24N) and 54 ribosomal protein overexpressing *E. coli* strains were spotted on LB agar medium containing 0.1 mM IPTG with tetracycline, streptomycin, or erythromycin. Colonies were counted after an overnight incubation. The number of colonies are presented in Figures S3 and S4. Correlation between the number of colonies formed in presence of Zn and antibiotics were analyzed.

### Overexpression of proteins other than ribosomal proteins does not lead to Zn resistance in *E. coli*

In subsequent analyses, we used strains overexpressing RpsO and RpsT as representative strains that overexpressed ribosomal proteins and showed Zn resistance. We next examined whether overexpression of proteins other than ribosomal proteins could also induce zinc resistance. The overexpression of GFPuv by isopropyl-β-D-thiogalactopyranoside (IPTG) did not induce Zn resistance (**Fig. 4A**). However, overexpression of the ribosomal proteins RpsO and RpsT by IPTG induced Zn resistance in *E. coli*, whereas Zn resistance was not observed when no IPTG was added, and RpsO and RpsT were not overexpressed (**Fig. 4A**). Since GFPuv is not an endogenous protein of *E. coli*, we examined whether the overexpression of *E. coli* proteins other than ribosomal proteins caused Zn resistance. Overexpression of proteins present in the cytosol (ThrA, Mog, CaiD) (18, 19, 20), membrane proteins (NhaA, KefC, SetA) (21), and complex-forming proteins (CarA, LeuD, IlvI) (22, 23, 24) by IPTG did not induce Zn resistance in *E. coli* (**Fig. 4B**). These results suggest that Zn resistance induced by overexpression is specific to ribosomal proteins.

**FIG 4.**
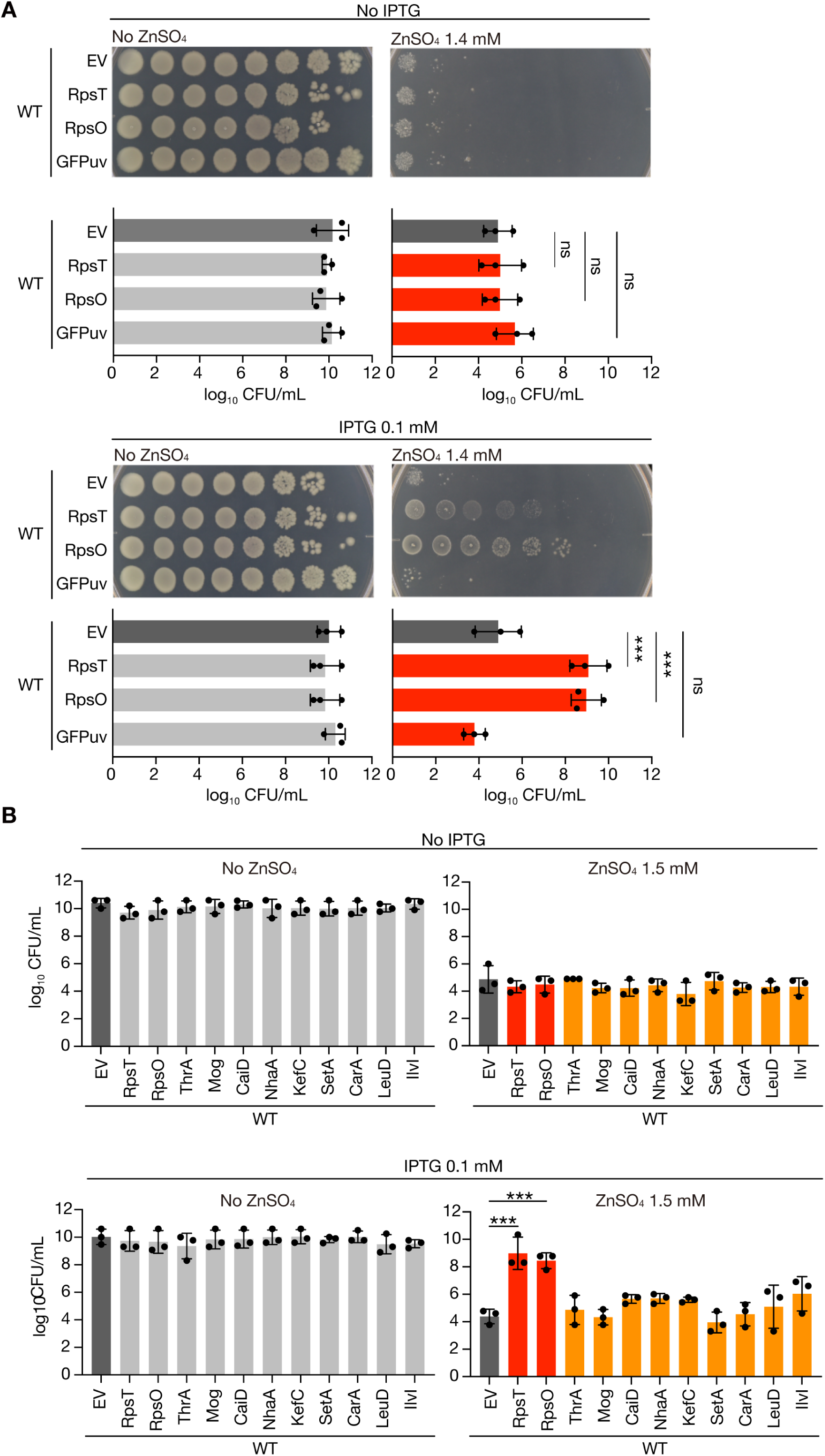
Overexpression of proteins other than ribosomal proteins does not lead to Zn resistance. A. Ten-fold serial dilutions of overnight cultures of an empty vector-transformed *E. coli* strain (WT/pCA24N), ribosomal protein-overexpressing strains (WT/pCA24N-RpsT, WT/pCA24N-RpsO), and GFPuv overexpressing strain (WT/pCA24N-GFPuv) were spotted on LB agar medium with or without 0.1 mM IPTG or 1.5 mM Zn, and colonies were counted after overnight incubation. Data represent mean ± SD of three independent experiments. \*\*\**p* < 0.001; ns, not significant [one-way analysis of variance (ANOVA) with Dunnett’s multiple comparison test]. B. Ten-fold serial dilutions of overnight cultures of an empty vector-transformed *E. coli* strain (WT/pCA24N), a ribosomal protein-overexpressing strain (WT/pCA24N-RpsT, WT/pCA24N-RpsO), and nine non-ribosomal protein-overexpressing strains were spotted on LB agar medium with or without 0.1 mM IPTG or 1.5 mM Zn. The colonies were counted after overnight incubation. Data represent mean ± SD of three independent experiments. \*\*\**p* < 0.001; ns, not significant (one-way ANOVA with Dunnett’s multiple comparison test).

### Zn resistance by overexpression of ribosomal proteins is ZntA-dependent

Zn accumulated in the cytoplasm is excreted by the ZntA. We investigated whether ZntA is required for Zn resistance caused by the overexpression of RpsO and RpsT. Because the *zntA*-knockout mutant was sensitive to Zn, the analysis was performed in the presence of 0.4 mM ZnSO_4_. Under 0.4 mM ZnSO_4_, no zinc resistance was found in the *zntA*-knockout mutant when RpsO and RpsT were overexpressed (**Fig. 5A**). Under 0.4 mM ZnSO_4_, the empty vector-transformed wild-type strain showed no growth sensitivity, and no Zn resistance was observed in the RpsO*-* and RpsT*-*overexpressing strains (**Fig. 5B**). In contrast, as mentioned above, at 1.4 mM ZnSO_4_ concentration, the strains overexpressing RpsO and RpsT were more resistant to Zn than the empty vector-transformed wild-type strain (**Fig. 2A**). Therefore, ZntA is required for Zn resistance through the overexpression of RpsO and RpsT.

**FIG 5.**
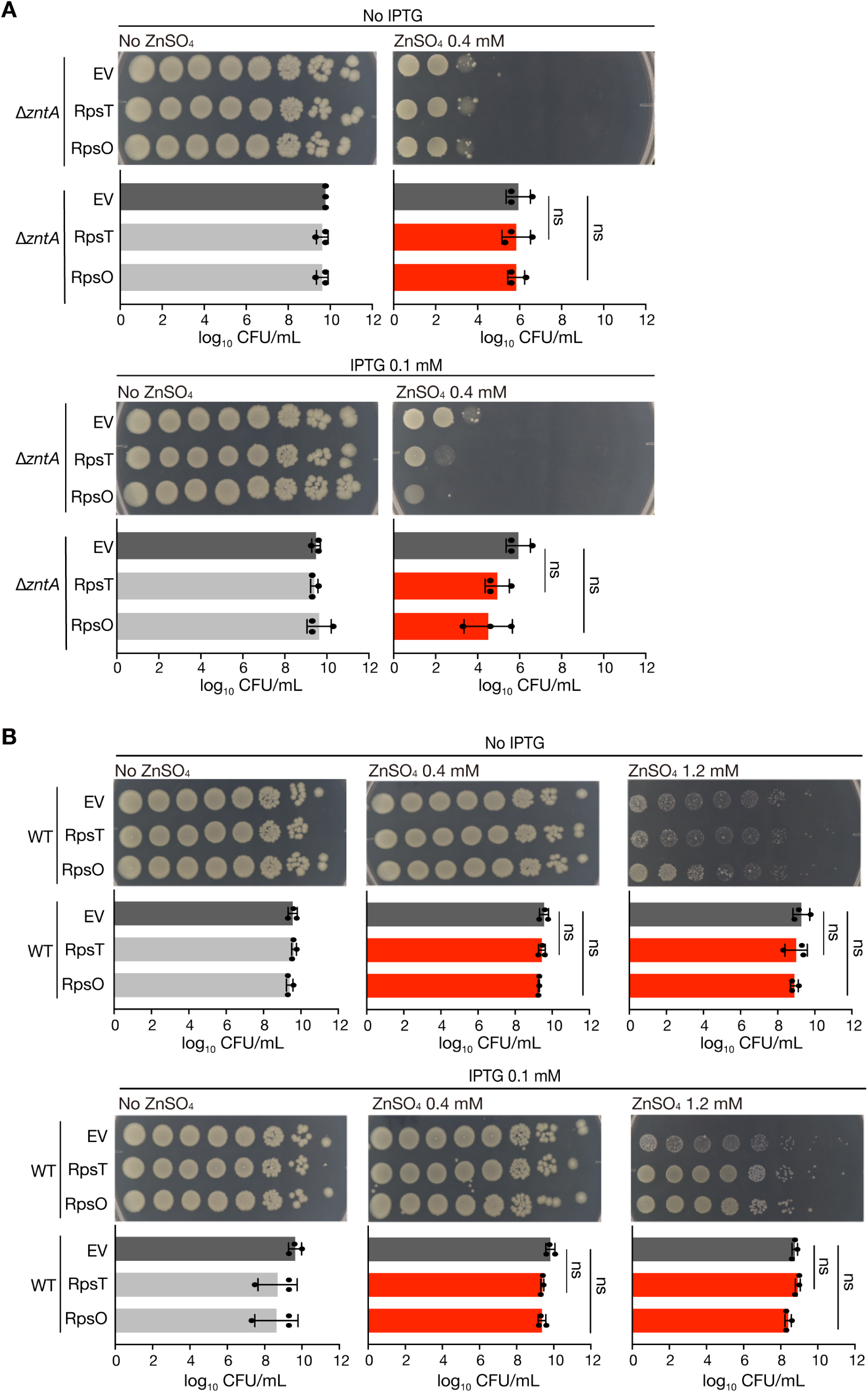
Knockout of *zntA* diminishes Zn resistance induced by the overexpression of ribosomal proteins. A. Ten-fold serial dilutions of overnight cultures of an empty vector-transformed *zntA*-knockout mutant (Δ*zntA*/pCA24N) and ribosomal protein-overexpressing *zntA*-knockout mutants (Δ*zntA*/pCA24N-RpsT, Δ*zntA*/pCA24N-RpsO) were spotted on LB agar medium with or without 0.1 mM IPTG or 1.2 mM Zn. Colonies were counted after overnight incubation. Data represent mean ± SD of three independent experiments. ns, not significant (one-way ANOVA with Dunnett’s multiple comparison test). B. Ten-fold serial dilutions of overnight cultures of an empty vector-transformed *E. coli* (WT/pCA24N), ribosomal protein-overexpressing strains (WT/pCA24N-RpsT, WT/pCA24N-RpsO) were spotted on LB agar medium containing IPTG or Zn at the indicated concentrations. Colonies were counted after overnight incubation. Data represent mean ± SD of three independent experiments. ns, not significant (one-way ANOVA with Dunnett’s multiple comparison test).

### Zn resistance by overexpression of ribosomal proteins is dependent on Lon protease

When bacteria are exposed to stress, such as nutrient starvation, excess intracellular ribosomes are degraded by Lon protease, which leads to stress tolerance (13). We investigated whether Lon protease was involved in the Zn resistance of *E. coli* induced by the overexpression of ribosomal proteins. Because the *lon*-knockout mutant was more sensitive to Zn than the wild-type strain, the analysis was performed in the presence of 1.2 mM ZnSO_4_. No Zn resistance was observed in the *lon*-knockout mutant when RpsO and RpsT were overexpressed (**Fig. 6A**). Therefore, the Lon protease is required for Zn resistance by the overexpression of RpsO and RpsT.

**FIG 6.**
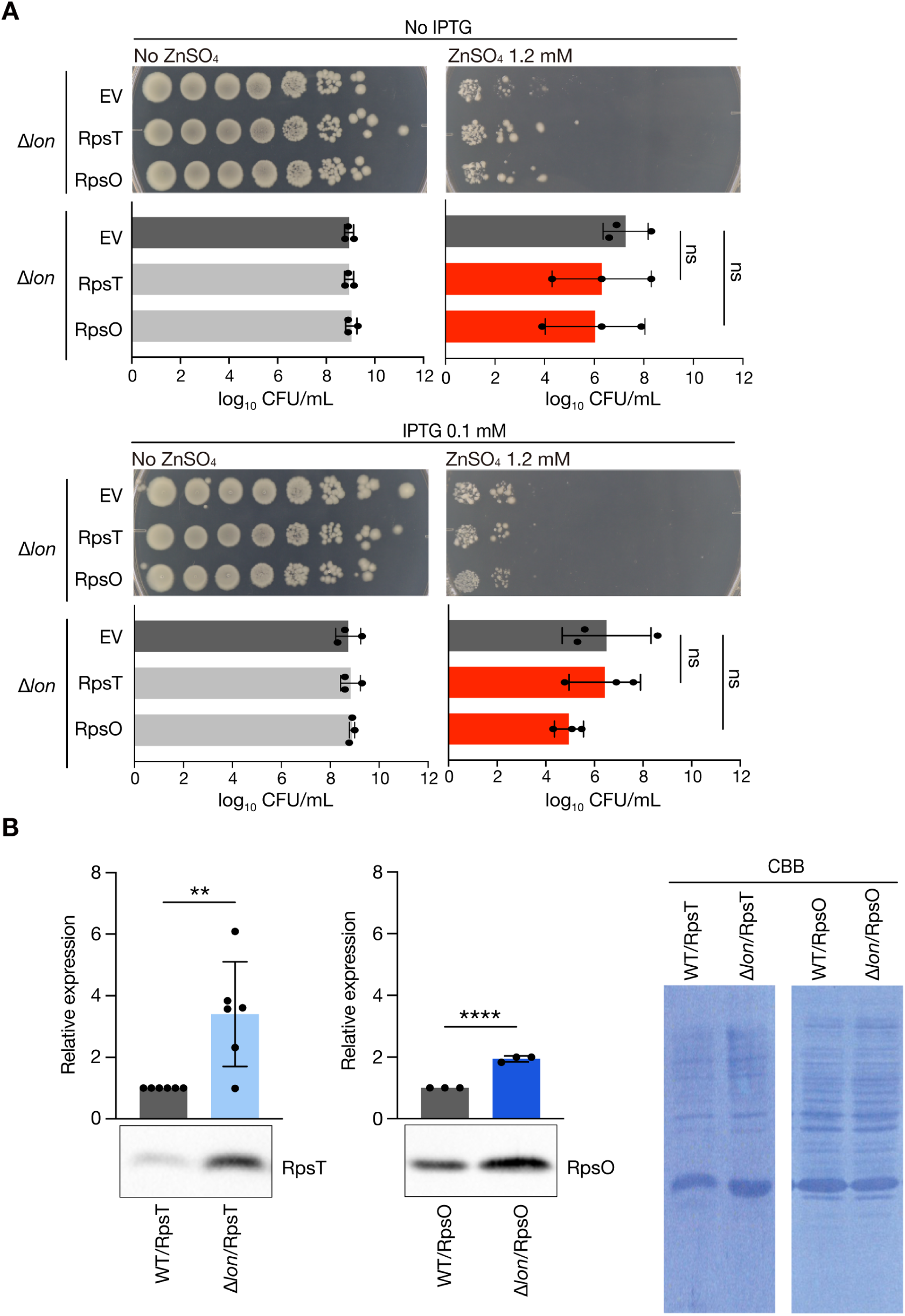
Knockout of *lon* diminishes Zn resistance caused by the overexpression of ribosomal proteins. A. Ten-fold step dilutions of overnight cultures of an empty vector-transformed *lon*-knockout mutant (Δ*lon*/pCA24N) and ribosomal protein-overexpressing *lon*-knockout mutants (Δ*lon*/pCA24N-RpsT, Δ*lon*/pCA24N-RpsO) were spotted on LB agar medium containing IPTG or Zn at the indicated concentrations. Colonies were counted after overnight incubation. Data represent mean ± SD of three independent experiments. ns, not significant (one-way ANOVA with Dunnett’s multiple comparison test). B. Overnight cultured cells of ribosomal protein-overexpressing wild-type *E. coli* (WT/pCA24N-RpsT, WT/pCA24N-RpsO) and ribosomal protein-overexpressing *lon*-knockout mutant (Δ*lon*/pCA24N-RpsT, Δ*lon*/pCA24N-RpsO) were collected and subjected to western blot analysis for detecting RpsT and RpsO expression. Data represent mean ± SD of three to six independent experiments. \**p* < 0.05; \*\*\*\**p* < 0.0001 (student *t*-test). Right image represents the membranes stained with Coomassie brilliant

We examined whether the overexpressed ribosomal proteins were degraded by Lon protease. The levels of RpsO or RpsT were higher in the *lon-*knockout mutant than in the wild-type strain (**Fig. 6B**). Total protein levels did not significantly change in the *lon*-knockout mutant compared to the wild-type strain (**Fig. 6B**). These results suggested that overexpressed RpsO and RpsT were degraded by Lon protease.

## Discussion

This study revealed that overexpression of 48 out of the 54 ribosomal proteins made *E. coli* resistant to Zn. Overexpression of proteins other than ribosomal proteins did not lead to Zn resistance. This is the first report to demonstrate that the overexpression of ribosomal proteins induces Zn resistance in *E. coli*.

In the present study, the strains overexpressing ribosomal proteins did not show resistance to metal ions other than Zn. This suggests that the overexpression of ribosomal proteins induces resistance specifically to Zn. The structural locations (25) and functions in the ribosome, and amino acid sequences were not conserved among the 48 ribosomal proteins that induced Zn resistance; therefore, the key characteristics of the ribosomal proteins that cause Zn resistance are unknown. We found a correlation between the degrees of resistance to streptomycin and Zn. Streptomycin inhibits the binding of N-formylmethionine-tRNA to the P site of ribosomes (26). Therefore, Zn may show toxicity by inhibiting the interaction between the ribosomal P site and N-formylmethionine-tRNA, similar to that by streptomycin, which may be affected by altered ribosome structure. Whether overexpressed ribosomal proteins alter the structure of ribosomal complex and interaction between ribosomes and tRNAs should be clarified in future.

Knockout of ZntA diminished Zn resistance induced by the overexpression of ribosomal proteins. This suggests that the overexpression of ribosomal proteins enhances ZntA activity through unknown pathways to cause Zn resistance. Furthermore, knockout of Lon protease (27), which is responsible for degrading aberrant proteins, abolished Zn resistance induced by the overexpression of ribosomal proteins. Since the accumulation of overexpressed ribosomal proteins was observed in the *lon*-knockout mutant, overexpressed ribosomal proteins were substrates for Lon protease, and ribosomal protein degradation by Lon protease, rather than ribosomal protein accumulation, may lead to Zn resistance. Ribosomal proteins are preferentially degraded by Lon protease during amino acid starvation and are used as amino acid sources (13). Because some amino acid-metabolizing enzymes have iron–sulfur clusters that are targets of Zn (28), excess Zn may inhibit amino acid-metabolizing enzymes and disrupt amino acid homeostasis in cells. Zn also inhibits leucine incorporation, leading to amino acid starvation (29). In these cases, ribosomal protein degradation by Lon protease may provide amino acids to suppress Zn-induced toxicity. In addition, Lon protease is involved in various bacterial physiological processes, such as pathogenicity, stress response, and quorum sensing, by regulating the amount of proteins that are direct targets for degradation as well as indirectly altering the transcription of diverse genes (30, 31, 32). For example, in *E. coli*, knockout of *lon* affects the expression of a large number of genes under the control of sigma S and GadE, which are involved in stress response and acid resistance (33). Based on these findings, it is possible that Lon protease is required for the expression of proteins involved in Zn resistance. Further analysis is needed to determine how Lon protease is involved in Zn resistance caused by the overexpression of ribosomal proteins.

This study demonstrates that the overexpression of ribosomal proteins induces ZntA- and Lon-mediated Zn resistance in *E. coli*. Further analyses are required to clarify the molecular mechanism of Zn resistance caused by the overexpression of ribosomal proteins and its physiological role.

## Materials and Methods

### Bacteria and culture conditions

*E. coli* BW25113 and its gene-knockout strains were cultured on Luria–Bertani (LB) agar medium, and single colonies were aerobically cultured at 37 °C in liquid LB medium. pCA24N-based plasmids-transformed *E. coli* was cultured on LB agar containing 30 µg/mL chloramphenicol, and pMW118-transformed *E. coli* was cultured on LB agar containing 100 µg/mL ampicillin. For overexpression of proteins, IPTG was added to the medium, and cells were cultured. Details of the bacteria and plasmids used in this study are listed in Table 1.

**Table 1.**
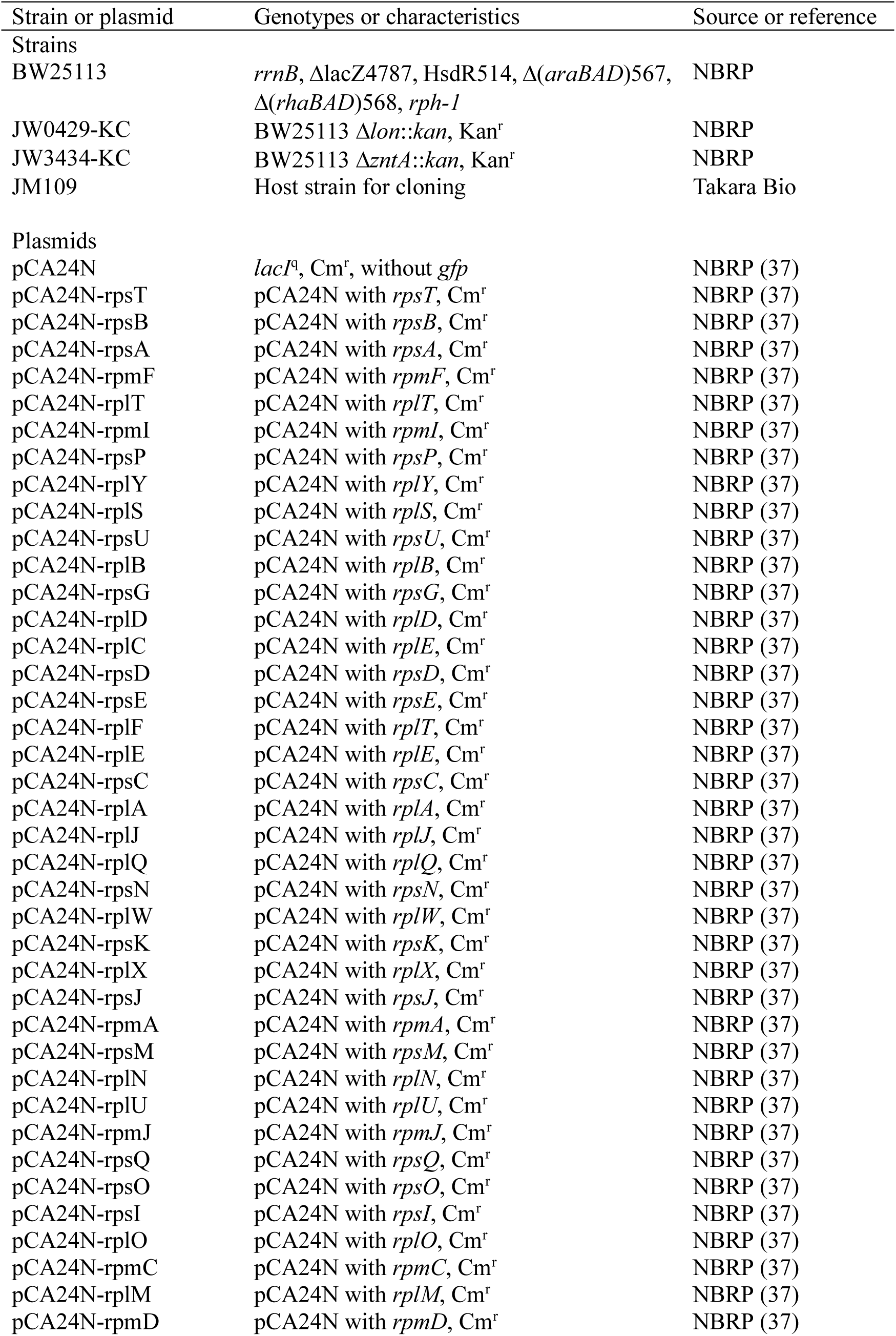
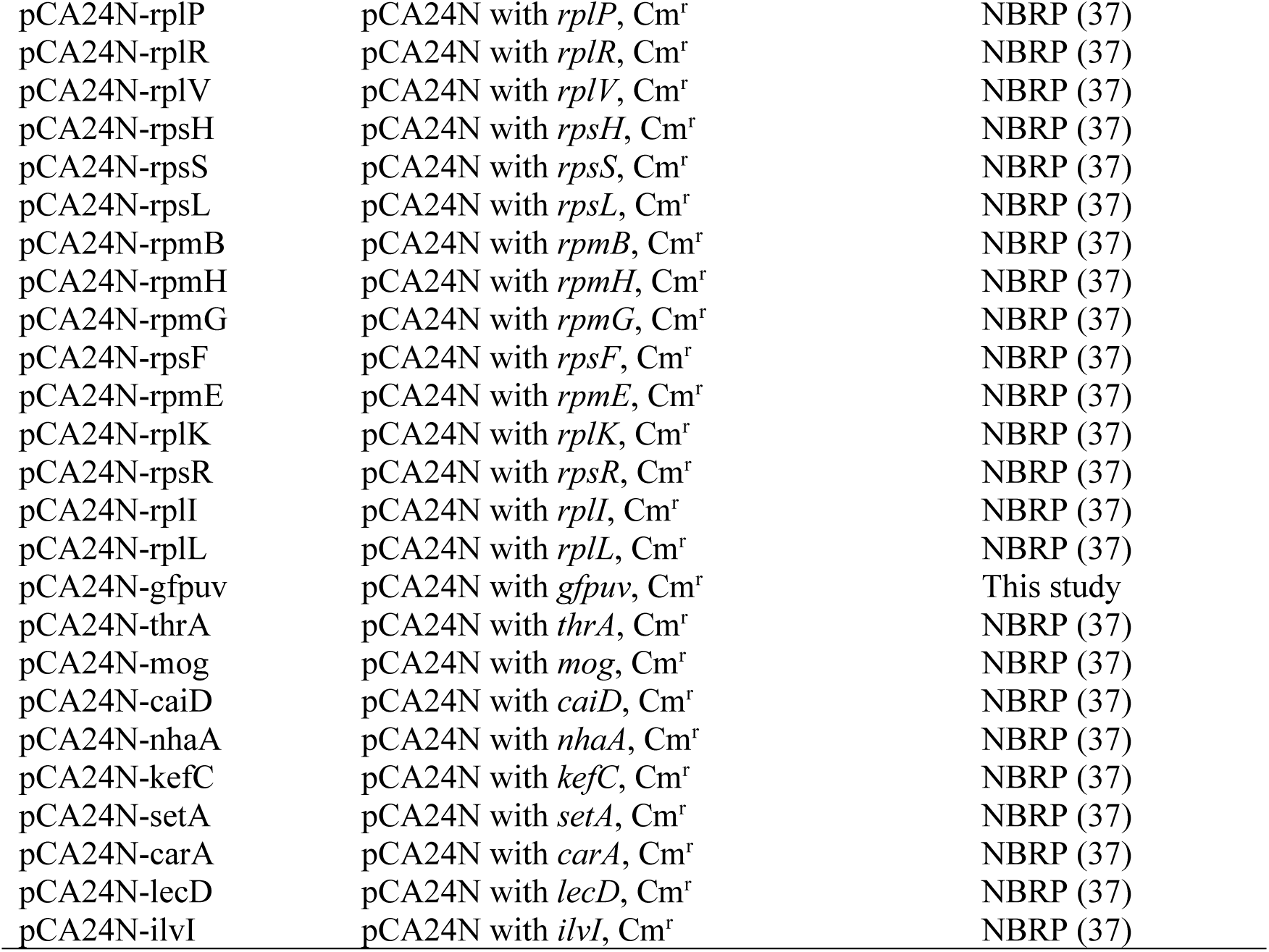
List of bacterial strains and plasmids used.

### Evaluation of bacterial resistance to metals and antimicrobial substances

To evaluate the resistance of the bacteria to Zn and antibiotics, autoclaved LB agar was mixed with 1.4 mM ZnSO_4_ (NACALAI TESQUE, Kyoto, Japan), 0.14 mM AgNO_3_ (NACALAI), 2.5 mM NiCl_2_ (NACALAI), 20 mM MnCl_2_ (NACALAI), 3.5 mM CuSO_4_ (Iwatsu Pharmaceuticals, Tokyo, Japan), 2.1 μg/mL tetracycline (Sigma-Aldrich, St. Louis, MO, USA), 12 μg/mL streptomycin (Sigma-Aldrich), or 140 μg/mL erythromycin (WAKO, Osaka, Japan) and poured into round or square petri dishes. Overnight cultures of *E. coli* were serially diluted by 10-fold in 96-well microplates, and 5 µL of the diluted culture was spotted on drug-containing LB agar medium. The plates were incubated overnight at 37 °C, and the colonies were photographed using a digital camera. The colonies were counted, and the CFU was calculated as previously described (34).

### Construction of gene-knockout mutant and transformation with plasmids

The *zntA* and *lon*-knockout mutants were constructed by transduction using P1 phage *vir* from the Keio collection as the phage donor and BW25113 as the recipient strain (35, 36). Plasmids overexpressing the 54 ribosomal proteins were extracted from *E. coli* strains carrying the ASKA library without *gfp* gene (37) and transformed into *E. coli* BW25113 cells. Colonies were grown on LB agar medium containing 30 µg/mL chloramphenicol.

To construct pCA24N carrying g*fpuv*, DNA fragments containing g*fpuv* was amplified by PCR using primer pairs (sfi1-GFP-F, 5’-CCGGCCCTGAGGGCCATGAGTAAAGGAGAAGAACT-3’; sfi1-GFP-R, 5’-GCGGCCGCATAGGCCTTATTTGTAGAGCTCATCCA-3’) and pGFPuv as a template. The amplified DNA fragment was inserted into SfiI site of pCA24N, resulting in pCA24N-gfpuv.

### Western blot analysis

*E. coli* strains were aerobically cultured in LB liquid medium at 37 °C, washed with phosphate-buffered saline, incubated with SDS sample buffer, and boiled for 5 min. The samples were electrophoresed on a 12.5% sodium dodecyl sulfate–polyacrylamide gel. Separated proteins were transferred onto polyvinylidene difluoride membranes (Immobilon P; Millipore). The membranes were treated with anti-6×His antibody diluted 1:5,000 in 1× Tris-buffered saline containing 0.05% Tween 20 (TBST). After three washes with TBST, the membranes were incubated with horseradish peroxidase-conjugated anti-mouse IgG (Promega, Japan) diluted 1:10,000 in 1× TBST. After three washes with TBST, the membranes were incubated with HRP substrate (Western Lighting Plus-ECL; PerkinElmer, MA, USA), and signals were detected using an iBright CL1500 Imaging System (Thermo Fisher Scientific, MA, USA).

### Statistics Analysis

All statistical analyses were performed using GraphPad PRISM software. Western blot results were analyzed using student’s *t*-test. Correlation was analyzed using Pearson’s correlation analysis. Spot assay results were analyzed using Dunnett’s test.

## Acknowledgements

This study was supported by Japan Society for the Promotion of Science (JSPS) Grants-in-Aid for Scientific Research (Grants 22K14892, 23K24131, 23K06130, 24K01760, 24K21872). This study was also supported by the Takeda Science Foundation (to C.K.), the Ichiro Kanehara Foundation (to C.K.), and the Ryobi Teien Memory Foundation (to K.I. and C.K.). We thank the National BioResource Project-E. coli (National Institute of Genetics, Japan) for providing the *E. coli* ASKA library and the Keio collection.

## References

[1] Rensing C, Helmann JD. 2017. Metal homeostasis and resistance in bacteria. Nat Rev Microbiol 15:338–350.

[2] Kasahara M, Anraku Y. 1974. Succinate- and NADH oxidase systems of Escherichia coli membrane vesicles. Mechanism of selective inhibition of the systems by zinc ions. J Biochem 76:967–976.

[3] Brazel EB, Tan A, Neville SL, Iverson AR, Udagedara SR, Cunningham BA, Sikanyika M, De Oliveira DMP, Keller B, Bohlmann L, El-Deeb IM, Ganio K, Eijkelkamp BA, McEwan AG, von Itzstein M, Maher MJ, Walker MJ, Rosch JW, McDevitt CA. 2022. Dysregulation of Streptococcus pneumoniae zinc homeostasis breaks ampicillin resistance in a pneumonia infection model. Cell Rep 38:110202.

[4] Li J, Ren X, Fan B, Huang Z, Wang W, Zhou H, Lou Z, Ding H, Lyu J, Tan G. 2019. Zinc Toxicity and Iron-Sulfur Cluster Biogenesis in Escherichia coli. Applied and Environmental Microbiology 85:e01967–18.

[5] Djoko KY, Ong CY, Walker MJ, McEwan AG. 2015. The Role of Copper and Zinc Toxicity in Innate Immune Defense against Bacterial Pathogens. J Biol Chem 290:18954–18961.

[6] Botella H, Peyron P, Levillain F, Poincloux R, Poquet Y, Brandli I, Wang C, Tailleux L, Tilleul S, Charrière GM, Waddell SJ, Foti M, Lugo-Villarino G, Gao Q, Maridonneau-Parini I, Butcher PD, Castagnoli PR, Gicquel B, de Chastellier C, Neyrolles O. 2011. Mycobacterial P1-Type ATPases Mediate Resistance to Zinc Poisoning in Human Macrophages. Cell Host Microbe 10:248–259.

[7] Barisch C, Kalinina V, Lefrançois LH, Appiah J, López-Jiménez AT, Soldati T. 2018. Localization of all four ZnT zinc transporters in Dictyostelium and impact of ZntA and ZntB knockout on bacteria killing. J Cell Sci 131:jcs222000.

[8] Rensing C, Mitra B, Rosen BP. 1997. The zntA gene of Escherichia coli encodes a Zn(II)-translocating P-type ATPase. Proc Natl Acad Sci U S A 94:14326–14331.

[9] Lee SM, Grass G, Haney CJ, Fan B, Rosen BP, Anton A, Nies DH, Rensing C. 2002. Functional analysis of the Escherichia coli zinc transporter ZitB. FEMS Microbiol Lett 215:273–278.

[10] Ramakrishnan V. 2002. Ribosome structure and the mechanism of translation. Cell 108:557–572.

[11] VanBogelen RA, Neidhardt FC. 1990. Ribosomes as sensors of heat and cold shock in Escherichia coli. Proc Natl Acad Sci U S A 87:5589–5593.

[12] Zhou X, Liao W-J, Liao J-M, Liao P, Lu H. 2015. Ribosomal proteins: functions beyond the ribosome. J Mol Cell Biol 7:92–104.

[13] Kuroda A, Nomura K, Ohtomo R, Kato J, Ikeda T, Takiguchi N, Ohtake H, Kornberg A. 2001. Role of inorganic polyphosphate in promoting ribosomal protein degradation by the Lon protease in E. coli. Science 293:705–708.

[14] Zhang S, Scott JM, Haldenwang WG. 2001. Loss of ribosomal protein L11 blocks stress activation of the Bacillus subtilis transcription factor sigma(B). J Bacteriol 183:2316–2321.

[15] Bursać S, Brdovčak MC, Pfannkuchen M, Orsolić I, Golomb L, Zhu Y, Katz C, Daftuar L, Grabušić K, Vukelić I, Filić V, Oren M, Prives C, Volarevic S. 2012. Mutual protection of ribosomal proteins L5 and L11 from degradation is essential for p53 activation upon ribosomal biogenesis stress. Proc Natl Acad Sci U S A 109:20467–20472.

[16] Shirakawa R, Ishikawa K, Furuta K, Kaito C. 2023. Knockout of ribosomal protein RpmJ leads to zinc resistance in Escherichia coli. PLOS ONE 18:e0277162.

[17] Kaczanowska M, Rydén-Aulin M. 2007. Ribosome biogenesis and the translation process in Escherichia coli. Microbiol Mol Biol Rev 71:477–494.

[18] Peabody MA, Laird MR, Vlasschaert C, Lo R, Brinkman FSL. 2016. PSORTdb: expanding the bacteria and archaea protein subcellular localization database to better reflect diversity in cell envelope structures. Nucleic Acids Res 44:D663–668.

[19] Ishihama Y, Schmidt T, Rappsilber J, Mann M, Hartl FU, Kerner MJ, Frishman D. 2008. Protein abundance profiling of the Escherichia coli cytosol. BMC Genomics 9:102.

[20] Díaz-Mejía JJ, Babu M, Emili A. 2009. Computational and experimental approaches to chart the Escherichia coli cell-envelope-associated proteome and interactome. FEMS Microbiol Rev 33:66–97.

[21] Daley DO, Rapp M, Granseth E, Melén K, Drew D, von Heijne G. 2005. Global topology analysis of the Escherichia coli inner membrane proteome. Science 308:1321–1323.

[22] Rubino SD, Nyunoya H, Lusty CJ. 1986. Catalytic domains of carbamyl phosphate synthetase. Glutamine-hydrolyzing site of Escherichia coli carbamyl phosphate synthetase. J Biol Chem 261:11320–11327.

[23] Fultz PN, Kwoh DY, Kemper J. 1979. Salmonella typhimurium newD and Escherichia coli leuC genes code for a functional isopropylmalate isomerase in Salmonella typhimurium-Escherichia coli hybrids. J Bacteriol 137:1253–1262.

[24] Gollop N, Damri B, Barak Z, Chipman DM. 1989. Kinetics and mechanism of acetohydroxy acid synthase isozyme III from Escherichia coli. Biochemistry 28:6310–6317.

[25] Gabashvili IS, Agrawal RK, Spahn CM, Grassucci RA, Svergun DI, Frank J, Penczek P. 2000. Solution structure of the E. coli 70S ribosome at 11.5 A resolution. Cell 100:537–549.

[26] Luzzatto L, Apirion D, Schlessinger D. 1968. Mechanism of action of streptomycin in E. coli: interruption of the ribosome cycle at the initiation of protein synthesis. Proc Natl Acad Sci U S A 60:873–880.

[27] Rosen R, Biran D, Gur E, Becher D, Hecker M, Ron EZ. 2002. Protein aggregation in Escherichia coli: role of proteases. FEMS Microbiol Lett 207:9–12.

[28] Lénon M, Arias-Cartín R, Barras F. 2022. The Fe-S proteome of Escherichia coli: prediction, function, and fate. Metallomics 14:mfac022.

[29] Anraku Y. 1968. Transport of sugars and amino acids in bacteria. 3. Studies on the restoration of active transport. J Biol Chem 243:3128–3135.

[30] Song X, Zhang H, Ma S, Song Y, Lv R, Liu X, Yang B, Huang D, Liu B, Jiang L. 2019. Transcriptome analysis of virulence gene regulation by the ATP-dependent Lon protease in Salmonella Typhimurium. Future Microbiol 14:1109–1122.

[31] Liu Y, Dong H, Peng X, Gao Q, Jiang H, Xu G, Qin Y, Niu J, Sun S, Li P, Ding J, Chen R. 2019. RNA-seq reveals the critical role of Lon protease in stress response and Brucella virulence. Microb Pathog 130:112–119.

[32] Yang N, Lan L. 2016. Pseudomonas aeruginosa Lon and ClpXP proteases: roles in linking carbon catabolite repression system with quorum-sensing system. Curr Genet 62:1–6.

[33] Heuveling J, Possling A, Hengge R. 2008. A role for Lon protease in the control of the acid resistance genes of Escherichia coli. Mol Microbiol 69:534–547.

[34] Uneme M, Ishikawa K, Furuta K, Yamashita A, Kaito C. 2024. Overexpression of the flagellar motor protein MotB sensitizes Bacillus subtilis to aminoglycosides in a motility-independent manner. PLoS One 19:e0300634.

[35] Baba T, Ara T, Hasegawa M, Takai Y, Okumura Y, Baba M, Datsenko KA, Tomita M, Wanner BL, Mori H. 2006. Construction of Escherichia coli K-12 in-frame, single-gene knockout mutants: the Keio collection. Mol Syst Biol 2:2006.0008.

[36] Yamamoto N, Nakahigashi K, Nakamichi T, Yoshino M, Takai Y, Touda Y, Furubayashi A, Kinjyo S, Dose H, Hasegawa M, Datsenko KA, Nakayashiki T, Tomita M, Wanner BL, Mori H. 2009. Update on the Keio collection of Escherichia coli single-gene deletion mutants. Mol Syst Biol 5:335.

[37] Kitagawa M, Ara T, Arifuzzaman M, Ioka-Nakamichi T, Inamoto E, Toyonaga H, Mori H. 2005. Complete set of ORF clones of Escherichia coli ASKA library (a complete set of E. coli K-12 ORF archive): unique resources for biological research. DNA Res 12:291–299.

